## Supplemental Material for "The level of HAND1 controls the specification of multipotent cardiac and extraembryonic progenitors"

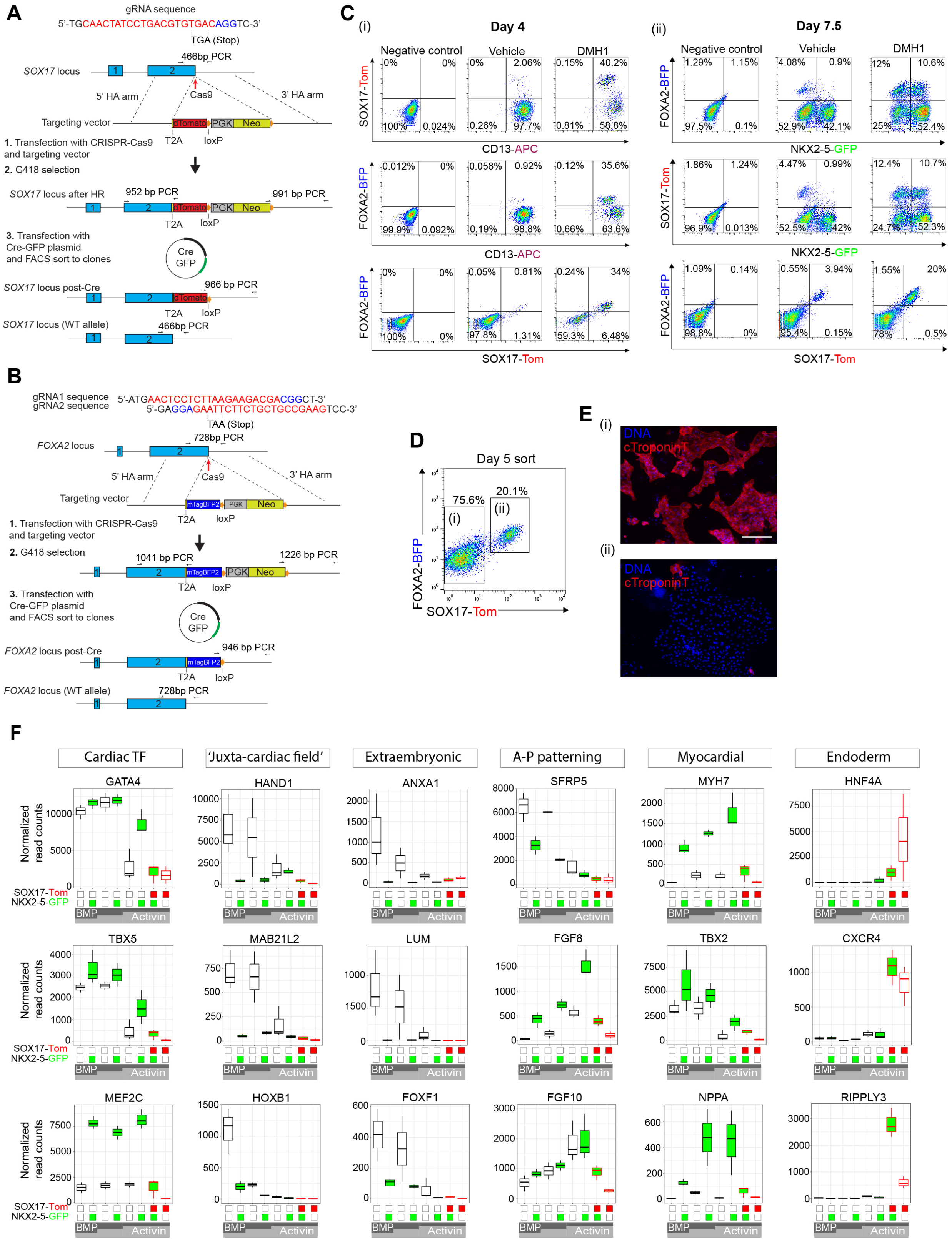

**Figure S1**. **Resolving cell diversity in mesendoderm differentiation**. A) Generation of *SOX17-T2A-dTomato* reporter knock-in hESCs by CRISPR-Cas9 gene targeting. A single gRNA was used to target the stop codon of *SOX17*. After homologous recombination, the integrated selection cassette was removed by Cre recombinase. One allele remained unedited. B)

Generation of *FOXA2-T2A-mTagBFP* reporter knock-in hESCs by CRISPR-Cas9 gene targeting. Two gRNAs were used to target the stop codon of *FOXA2*. After homologous recombination, the integrated selection cassette was removed by Cre recombinase. One allele remained unedited. C) Flow cytometric analyses at day 4 (i) and day 7.5 (ii) of differentiation of *FOXA2-BFP SOX17-Tom NKX2-5-GFP* triple reporter hESCs in control and DMH1-treated (day 2−3) conditions. The surface marker CD13-APC was included in the analysis at day 4. D) Flow cytometric sorting at day 5 by FOXA2-BFP and SOX17-Tomato into double negative (i) and double positive (ii) populations for differentiation. E) Immunostaining of cTroponinT from the sorted populations as in D. F) Gene expression by RNA-seq of selected markers in the 8 sorted populations based on SOX17-Tom and NKX2-5-GFP at day 7.5. The boxplots follow standard Tukey representations and are colored by the lineage markers. Scale bars represent 150 μm. A-P, anterior-posterior.

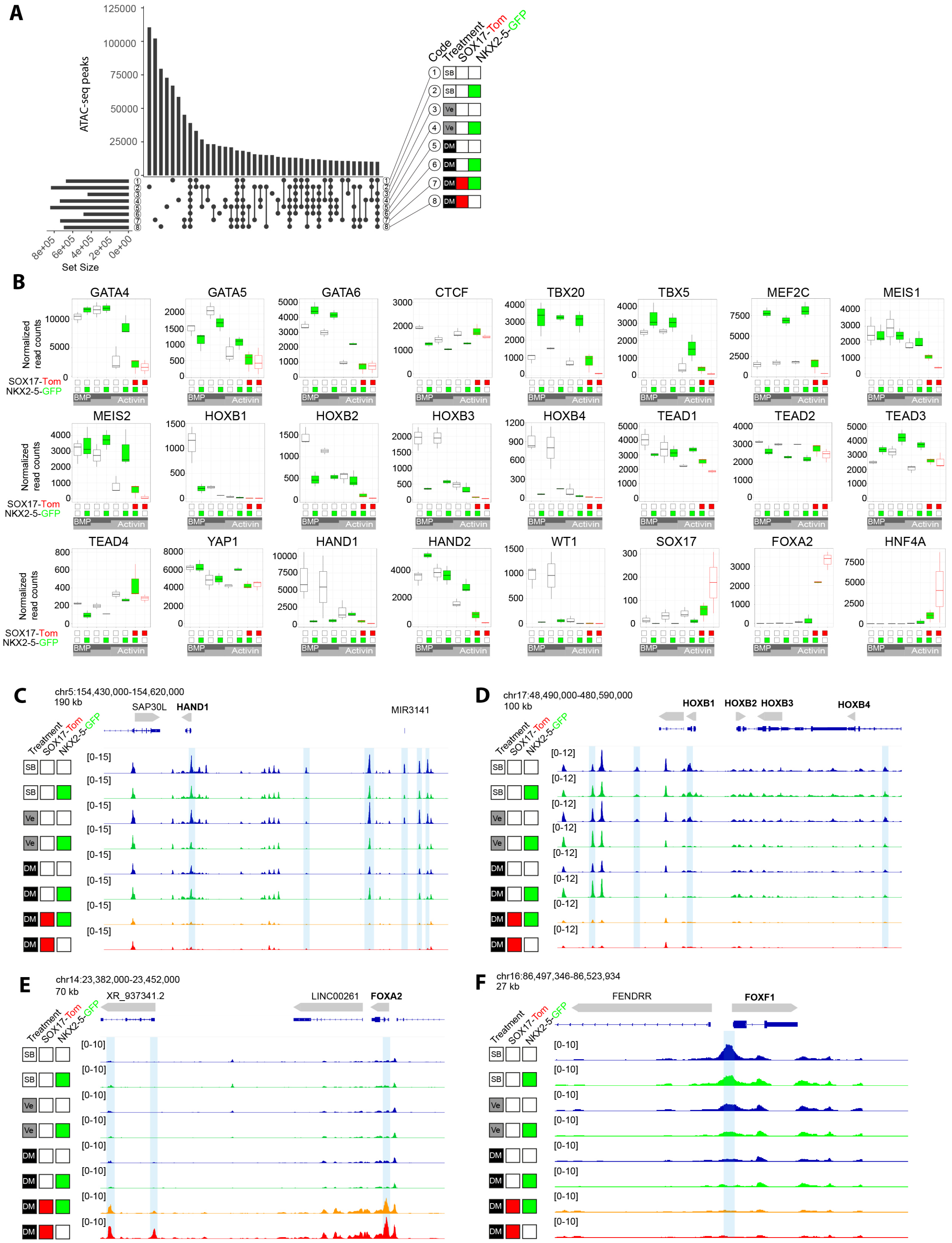


**Figure S2**. **ATAC-seq and transcription factor motif enrichment analysis with differentiation**. A) Upset plot of ATAC-seq peaks from the 8 sorted populations sorted based on SOX17-Tomato and NKX2-5-GFP at day 7.5 (number coded based on identity as indicated). Connections indicate common peak sets. B) Gene expression by RNA-seq of transcription factors relevant to the motif enrichment analysis. Normalized read counts are shown. The boxplots follow standard Tukey representations and are colored by the lineage markers. C–E) ATAC-seq tracks colored according to marker identity around C) *HAND1*, D) *HOXB1–4*, E) *FOXA2* and F) *FOXF1* loci (RNA expression of these genes is shown in B or **Figure S1F**. Some differentially accessible regions are highlighted in blue. Genomic coordinates are shown top left.

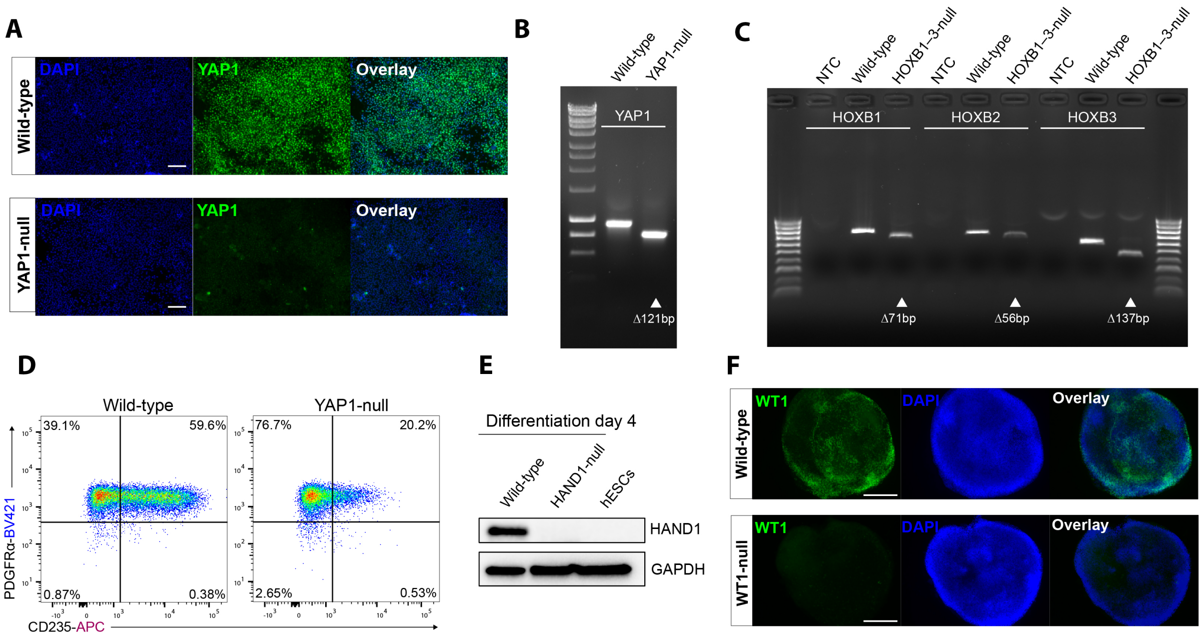


**Figure S3. Validation of hESC gene knockouts by CRISPR-Cas9**. A) Immunostaining of YAP1 in wild-type and YAP1-null hESCs after CRISPR-Cas9 mutagenesis. B) PCR showing expected 121bp frameshift-causing deletion in *YAP1* in YAP1-null hESCs. C) PCR showing expected 71bp, 56bp, and 137bp frameshift-causing deletions in *HOXB1*, *HOXB2*, and *HOXB3* respectively in HOXB1–3-null hESCs. D) Flow cytometric analysis of PDGFRα and CD235 in day 4 EBs derived from wild-type and YAP1-null hESCs. E) HAND1 western blot in day 4 EBs derived from wild-type and HAND1-null hESCs. Undifferentiated wild-type hESCs are shown as a negative control. F) Immunostaining of WT1 in wild-type and WT1-null day 10 EBs. Scale bars represent 150 $\mu$m in A and 300 $\mu$m in F. NTC, no template control.

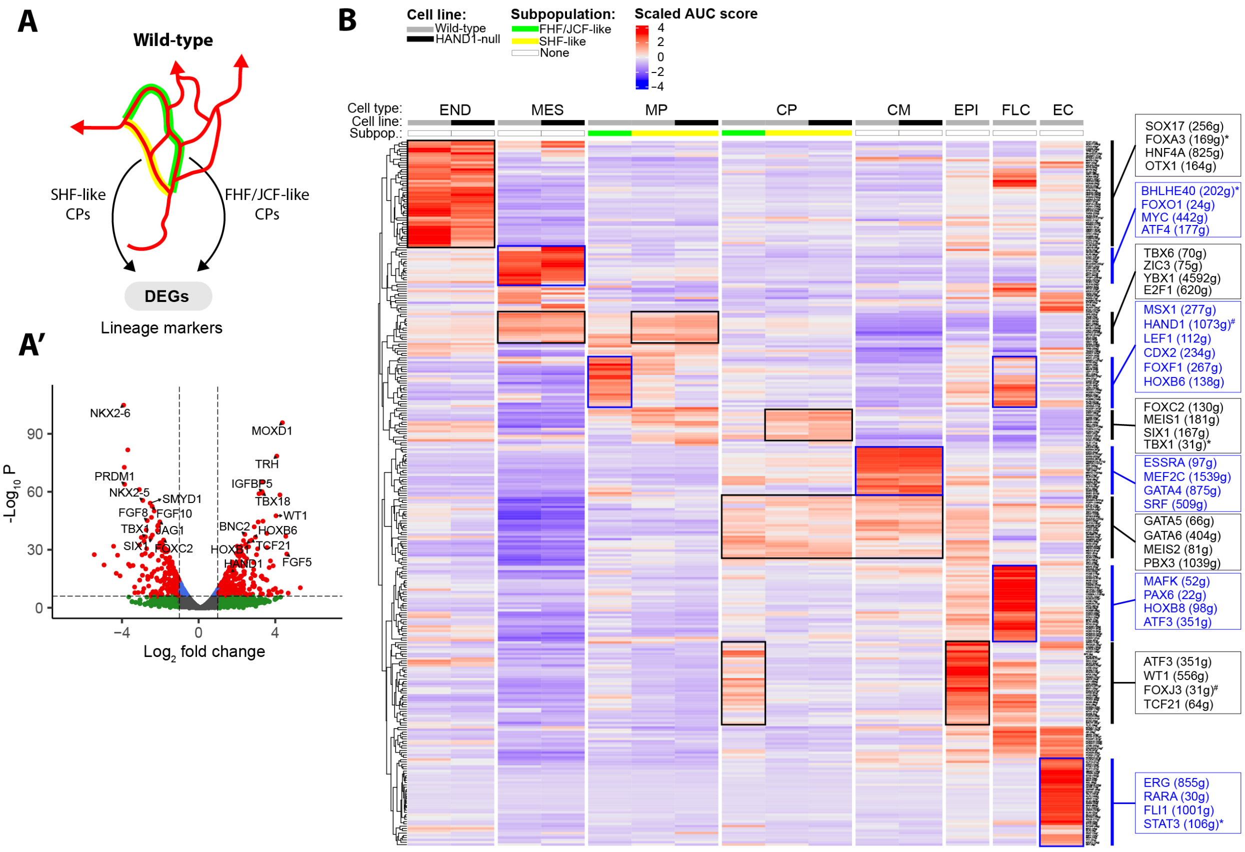


**Figure S4**. **Gene expression and regulatory network analysis during differentiation**. A) Differential gene expression of cardiac progenitors extracted from the two lineage trajectories of the wild-type differentiation. A') A volcano plot displaying the results of the DE analysis (FHF/JCF-like vs SHF-like) with some significant genes highlighted. B) Heatmap and hierarchical clustering showing the scaled target gene AUC score of the 307 regulons detected by SCENIC by cell line, cell type, and subpopulation. Clustering is by rows (regulons). AUC values were converted to a z-score and centered on the mean. Select markers present in clusters are highlighted. *Identified in HAND1-null population. #Identified in CPC lineage analysis. AUC, area under the curve; CPC, cardiac progenitor cell; CM, cardiomyocyte; EC, endothelial cells; END, endoderm; EPI, epicardial; FHF, first heart field; FLC, fibroblast-like cells; JCF, juxta-cardiac field; MES, mesoderm; MP, mesodermal progenitors; SCENIC, single-cell regulatory network interference and clustering; SHF, second heart field.

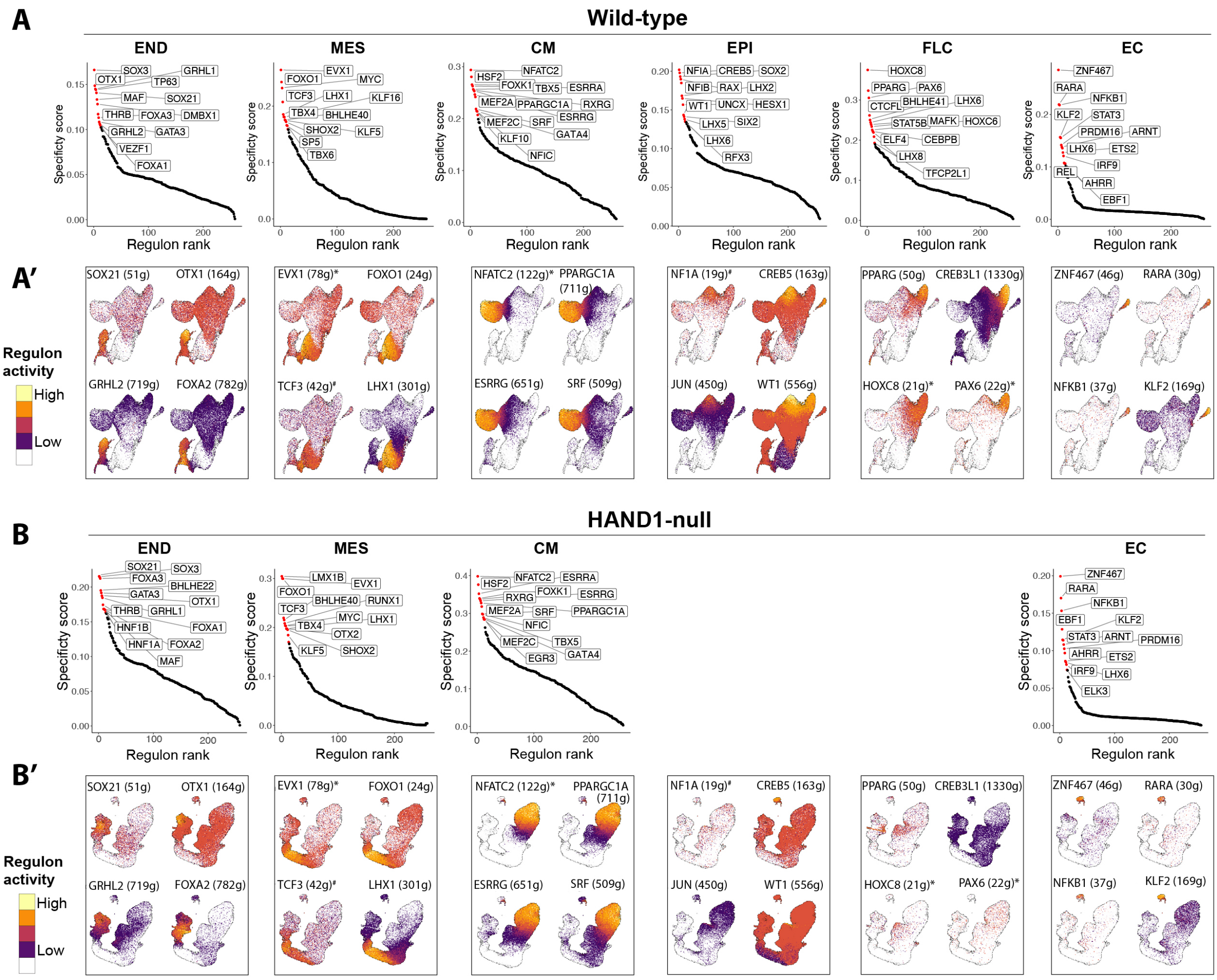


**Figure S5**. **SCENIC regulon specificity ranking in different populations during differentiation**. A) Regulon specificity scores ranked for different cell types in wild-type and B) in HAND1-null populations. The HAND1-null line did not generate EPI or FLC types. The top 5% are highlighted red and several are labelled. A') UMAP plots showing the activity of some key cell type-biased regulons in wild-type and in B') HAND1-null cells. *Identified in HAND1-null population. #Identified in CPC lineage analysis. AUC, area under the curve; CM, cardiomyocyte; EC, endothelial cells; END, endoderm; EPI, epicardial; MES, mesoderm; SCENIC, single-cell regulatory network interference and clustering.

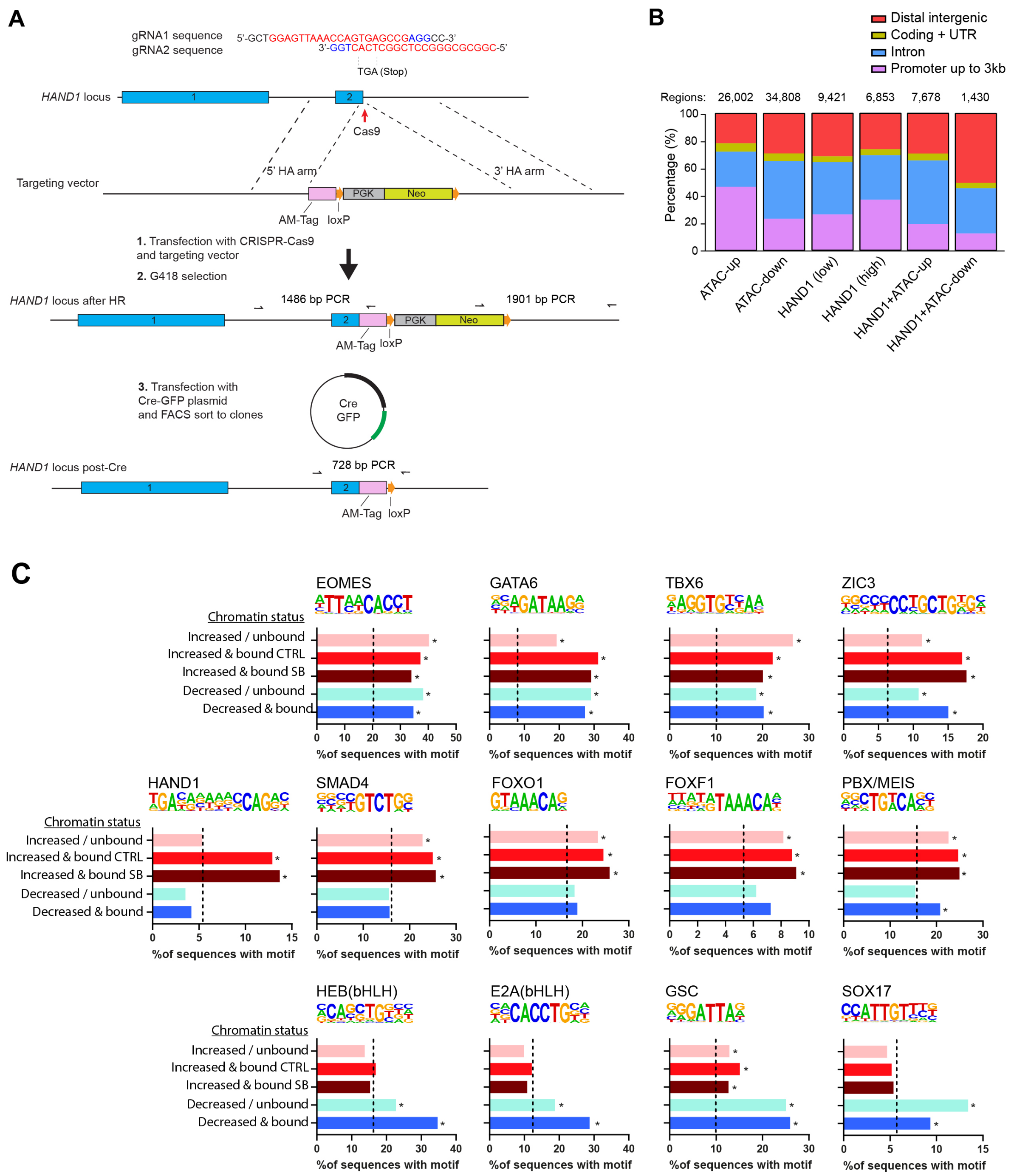


**Figure S6. The impact of HAND1 on chromatin in mesoderm**. A) Generation of *HAND1-AM-Tag* (Active Motif) knock-in hESCs by CRISPR-Cas9 gene targeting. Two gRNAs were used to target the stop codon of *HAND1*. After homologous recombination, the integrated selection cassette was removed by Cre recombinase. The knock-in was biallelic. B) Annotation of ATAC-seq (increased or decreased accessibility by HAND1 and HAND1 ChIP-seq peaks (all peaks or intersected with the ATAC peaks). C) Motif enrichment analysis of the regions indicated in **Figure 5C'**. The percentage of sequences with each motif is shown. * Significance above background (p<1E-6). UTR, untranslated region.

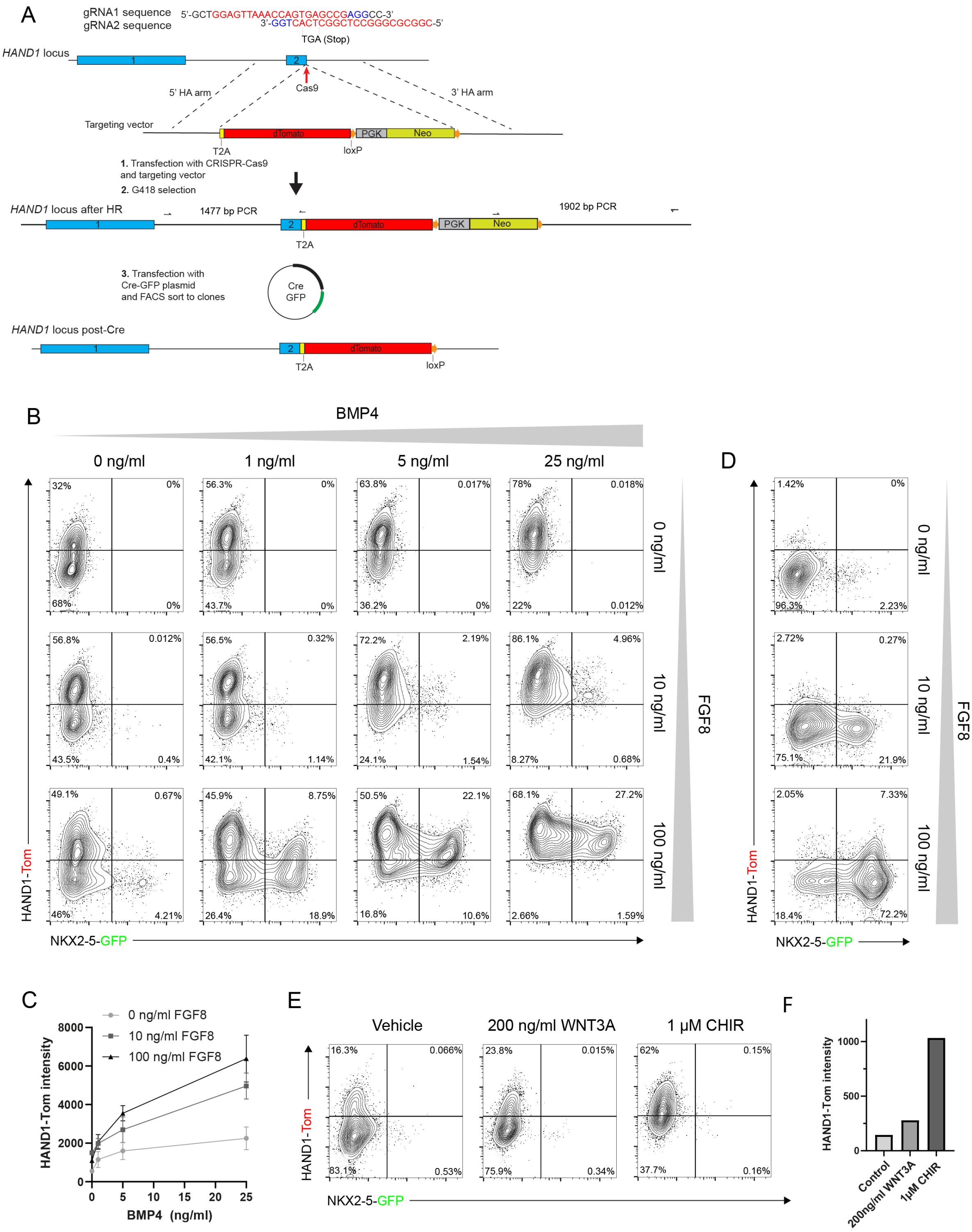


**Figure S7**. **BMP, FGF and WNT signaling pathways regulate the expression of *HAND1***. A) Generation of *HAND1-T2A-dTomato* reporter knock-in hESCs by CRISPR-Cas9 gene targeting. Two gRNAs were used to target the stop codon of HAND1. After homologous recombination, the integrated selection cassette was removed by Cre recombinase. The knock-in was biallelic. B) Flow cytometric analysis of HAND1-Tom and NKX2-5-GFP in progenitor cells derived from SB-treated EBs and cultured in different concentrations of BMP4 and FGF8. C) Median fluorescence intensity values of HAND1-Tomato with FGF8 and BMP4 exposure. D) Flow cytometric analysis of HAND1-Tom and NKX2-5-GFP in progenitor cells derived from DMH1-treated EBs and cultured in different concentrations of FGF8. E) Flow cytometric analysis of HAND1-Tom and NKX2-5-GFP in progenitor cells derived from SB-treated EBs and cultured in WNT3A or CHIR99021. F) Median fluorescence intensity values of HAND1-Tom with WNT3A and CHIR exposure. Data in C represent mean ± SD, n = 3 independent biological experiments. EB, embryoid body.

**Table S1. gRNAs sequences for fluorescent reporter knock-ins**

| gRNA | Sequence 5’-3’ |
|---|---|
| SOX17 KI 1 | CAACTATCCTGACGTGTGAC |
| FOXA2 KI 1 | AACTCCTCTTAAGAAGACGA |
| FOXA2 KI 2 | GAAGCCGTCGTCTTCTTAAG |
| HAND1 KI 1 | GGAGTTAAACCAGTGAGCCG |
| HAND1 KI 2 | CGGCGCGGGCCTCGGCTCAC |

**Table S2. gRNA sequences for gene knockouts**

| gRNA | Sequence 5’-3’ |
|---|---|
| HAND1 KO 1 | AGCGCGAGGCCGGACCGAAG |
| HAND1 KO 2 | CGCTTGGCGGCCGTCTTGGC |
| WT1 KO 1 | CGCTCCCGCAGGTTACAGCA |
| WT1 KO 2 | GATCCTCATGCTTGAATGAG |
| HOXB1 KO 1 | ACAGAGTGGGTACTCTAAGA |
| HOXB1 KO 2 | ACCGCCTGAGCCGAGCTTGG |
| HOXB2 KO 1 | CACCAGCCTCCGGCAGTCCC |
| HOXB2 KO 2 | CAGTTCCAGCAGCTGCGTGT |
| HOXB3 KO 1 | GAGGGGACAAGAGCCCCCCG |
| HOXB3 KO 2 | TCTTGATCTGCCGCTCGCTG |
| YAP1 KO 1 | GCTGCGAAGGCGGCTGCCCT |
| YAP1 KO 2 | ATCAGATCGTGCACGTCCGC |

**Table S3. Antibodies**

| Antibody | Supplier | Identifier |
|---|---|---|
| Polyclonal Goat anti-HAND1 | R & D Systems | #AF3168 |
| Monoclonal Mouse anti-TNNT2 | BD Biosciences | #564767 |
| Monoclonal Mouse anti-PDGFRα | BD Biosciences | #528238 |
| Monoclonal Mouse anti-TRA-1-81-FITC | BD Biosciences | #560883 |
| Monoclonal Mouse anti-CD235a-APC | BD Biosciences | #561775 |
| Monoclonal Mouse anti-ACTN2 | Sigma-Aldrich | #A7811 |
| Monoclonal Mouse anti-WT1 | Abcam | #ab89901 |
| Polyclonal Rabbit anti-CD31 | ThermoFisher | #BMS137 |
| Polyclonal Rabbit anti-FOXA2 | Abcam | #ab40874 |
| Monoclonal Mouse anti-CD13-APC | BioLegend | #301705 |
| Monoclonal Mouse anti-Cardiac Troponin T | Proteintech | #15513-1-AP |

| | | |
|---|---|---|
| Monoclonal Rabbit anti-YAP | Abcam | #ab205270 |
| Mouse AbFlex® AM-Tag | Active Motif | #91111 |
| Alexa Fluor-488 Donkey Anti-Rabbit IgG | Invitrogen | #A21206 |
| Alexa Fluor-555 Donkey Anti-Mouse IgG | Invitrogen | #A31570 |
| Alexa Fluor-594 Donkey Anti-Mouse IgG | Invitrogen | #A21203 |
| Monoclonal Rabbit anti-SMAD1 | Cell Signalling Technology | #6944 |
| Monoclonal Rabbit anti-Phospho-SMAD1/5/9 | Cell Signalling Technology | #13820 |
| Monoclonal Rabbit anti-SMAD2 | Cell Signalling Technology | #5339 |
| Monoclonal Rabbit anti-Phospho-SMAD2 | Cell Signalling Technology | #3108 |
| HRP-linked anti-Rabbit IgG | Cell Signalling Technology | #7074 |

**Table S4 RT-qPCR Primers**

| Gene | Forward 5'-3' | Reverse 5'-3' |
|---|---|---|
| FOXA2 | GCTGGTCGTTTGTTGTGGC | TTCATGCCGTTCATCCCCAG |
| GUSB | CCACCTAGAATCTGCTGGCTAC | GTGCCCGTAGTCGTGATACCAA |
| HOXB1 | TTCAGCAGAACTCCGGCTAT | CCTCCGTCTCCTTCTGATTG |
| MESP1 | CTCTGTTGGAGACCTGGATG | CCTGCTTGCCTCAAAGTG |
| MIXL1 | GGTACCCCGACATCCACTT | GAGACTTGGCACGCCTGT |
| RPLPO | CACCATTGAAATCCTGAGTGATGT | TGACCAGCCCAAAGGAGAAG |
| SOX2 | CCCAGCAGACTTCACATGT | CCTCCCATTTCCCTCGTTTT |
| TBXT | ATCACCAGCCACTGCTTC | GGGTTCCTCCATCATCTCTT |
| TNNT2 | TTCGACCTGCAGGAGAAGTT | GCGGGTCTTGGAGACTTTCT |
| WT1 | TATTCTGTATTGGGCTCCGC | CAGCTTGAATGCATGACCTG |

**Table S5. Barcoding primers for ATAC-Seq/ChIPmentation**

| Barcoding Primer | Sequence 5'-3' |
|---|---|
| N701 | CAAGCAGAAGACGGCATACGAGATTCGCCTTAGTCTCGTGGGCTCGGAGATGT |
| N702 | CAAGCAGAAGACGGCATACGAGATCTAGTACGGTCTCGTGGGCTCGGAGATGT |
| N703 | CAAGCAGAAGACGGCATACGAGATTTCTGCCTGTCTCGTGGGCTCGGAGATGT |
| N704 | CAAGCAGAAGACGGCATACGAGATGCTCAGGAGTCTCGTGGGCTCGGAGATGT |
| N501 | AATGATACGGCGACCACCGAGATCTACACTAGATCGCTCGTCGGCAGCGTC |
| N502 | AATGATACGGCGACCACCGAGATCTACACCTCTCTATTCGTCGGCAGCGTC |
| N503 | AATGATACGGCGACCACCGAGATCTACACTATCCTCTTCGTCGGCAGCGTC |
| N504 | AATGATACGGCGACCACCGAGATCTACACAGAGTAGATCGTCGGCAGCGTC |
